# Cardiomyocyte Perm1 sustains cardiac bioenergetic reserve and exercise capacity

**DOI:** 10.64898/2026.10.03.753324

**Authors:** Shizuko Tachibana, Tina Le, Simon Schenk, Wei Feng, Ju Chen, Robert S. Ross, Stephan Lange, Yoshitake Cho

## Abstract

**Objective:** The PGC-1/ERR axis controls both steady-state mitochondrial respiration and the capacity to expand ATP production on demand, but whether this reserve function requires dedicated downstream effectors, and whether its loss is detectable at rest, remain unclear. We tested whether Perm1, a striated muscle-specific PGC-1/ERR-induced regulator, is required for cardiac bioenergetic reserve.

**Methods:** We generated cardiomyocyte-specific Perm1 knockout mice using constitutive αMHC-nCre and tamoxifen-inducible αMHC-MerCreMer drivers and followed to 12 months and to 4 months after tamoxifen, respectively. Mitochondrial respiration was measured by high-resolution respirometry and ultrastructure by electron microscopy; energetic capacity under load was tested by treadmill exercise with tissue ATP quantification and by β-adrenergic stimulation of neonatal cardiomyocytes.

**Results:** Perm1 deletion did not perturb cardiac development; neither constitutive nor adult-onset deletion impaired cardiac structure and systolic function through 12 months. Despite this, mitochondrial respiration was reduced by 20–25% without a change in total mitochondrial content or OxPhos subunit abundance. MICOS components MIC60 and MIC19 were selectively reduced, and mitochondria redistributed from subsarcolemmal to interfibrillar compartments. Perm1-deficient mice of both sexes showed ∼50% lower treadmill endurance at 6 and 12 months and failed to maintain cardiac ATP at a submaximal workload, whereas gastrocnemius ATP and plasma lactate were unaffected. β-Adrenergic stimulation raised ATP content in control cardiomyocytes, and Perm1 knockdown blunted this response.

**Conclusions:** Perm1 is a cardiomyocyte-intrinsic determinant of cardiac bioenergetic reserve: dispensable for resting function but required to expand ATP production when demand rises. This regulation is invisible to resting phenotyping and emerges only under stress.

## 1. Introduction

Mitochondrial oxidative capacity is not a fixed tissue property but a dynamically regulated one, adjusted to match energy demand. In striated muscle, this adjustment is governed largely by the PGC-1 coactivators and the estrogen-related receptors (ERRs), which together drive programs of mitochondrial biogenesis, substrate oxidation, and respiratory chain assembly [1–3]. Loss of this axis compromises not only steady-state oxidative metabolism but, more selectively, the capacity to expand energy production when demand rises. Brown adipocytes lacking all three ERRs, for example, maintain normal basal respiration yet fail to increase maximal respiration or spare respiratory capacity in response to adrenergic stimulation [4]. Whether an analogous reserve function operates in striated muscle, and which downstream effectors execute it, remains poorly defined. Combined loss of the upstream coactivators does not resolve this question: cardiac deletion of both PGC-1α and PGC-1β arrests mitochondrial biogenesis and causes perinatal heart failure [5], removing baseline oxidative capacity rather than selectively removing reserve.

The heart is the tissue in which energetic reserve is extremely important. Cardiomyocytes sustain continuous contraction while maintaining a remarkably small ATP pool and must increase ATP production several-fold within seconds when workload rises during exercise or sympathetic activation. The creatine kinase system keeps cardiac ATP concentration nearly constant across workloads by buffering local ATP and delaying any fall until mitochondrial production can accelerate through calcium-dependent activation of matrix dehydrogenases [6–8]. Because this buffering is finite, a defect confined to the capacity to accelerate oxidative phosphorylation (OxPhos) may leave resting cardiac function and resting ATP levels intact but impair them when ATP demand increases, such as with exercise.

We previously identified Perm1 (PGC-1 and ERR-induced regulator, muscle 1) as a striated muscle-specific, PGC-1/ERR-induced regulator of mitochondrial biogenesis and oxidative metabolism [9; 10]. Cardiomyocyte Perm1 expression rises progressively from early embryonic stages through adulthood, paralleling that of PGC-1 and ERR family members, and cardiac Perm1 overexpression enhances oxidative metabolism and protects against pressure overload-induced dysfunction [11–13]. These gain-of-function data establish Perm1 as sufficient to augment cardiac oxidative capacity, but do not elucidate the importance of endogenous Perm1, be it to cardiac development or function. Indeed, previous studies have shown that global Perm1 knockout (KO) downregulates oxidative and mitochondrial gene programs in the heart [14] yet its consequences for contractile function remain disputed; some studies report reduced systolic function [14; 15], whereas others find no detectable change [16; 17]. A likely reason for these conflicting findings is the direct and indirect confounding effects of global KO of Perm1 in non-cardiac tissues, such as skeletal muscle. Thus, the specific role of Perm1 in cardiac function remains to be fully defined.

To more specifically assess the role of Perm1 in the myocardium, we generated a cardiomyocyte-specific Perm1 model. We used αMHC-nCre and tamoxifen-inducible αMHC-MerCreMer mice, coupled with a floxed Perm1 model, to evaluate both developmental consequences of Perm1 loss and postnatal compensation, and we assessed cardiac structure, function, mitochondrial architecture, and energetics at rest and under defined physiological load. We find that cardiomyocyte Perm1 is not required for cardiac development or resting cardiac performance through 12 months of age. However, Perm1 is required to sustain mitochondrial respiratory capacity, maintain exercise capacity, and cardiac ATP during exercise. These findings define Perm1 as a cardiomyocyte-intrinsic determinant of bioenergetic reserve and illustrate a mode of metabolic regulation that is invisible to resting phenotyping.

## 2. Results

### Generation of Perm1 floxed mice and cardiomyocyte-specific Perm1 knockout line

To generate a conditional Perm1 allele, we inserted loxP sites flanking exon 2 of Perm1 using CRISPR/Cas9-mediated genome editing (**Fig. 1A**) [18]. PCR genotyping confirmed successful loxP insertion, with the floxed allele producing a diagnostic fragment approximately 40 bp larger than the wild-type band (∼200 bp) (**Fig. 1A and 1B**). Genomic sequencing verified that the two loxP sites were integrated in cis on the same chromosome.

**Figure 1.**
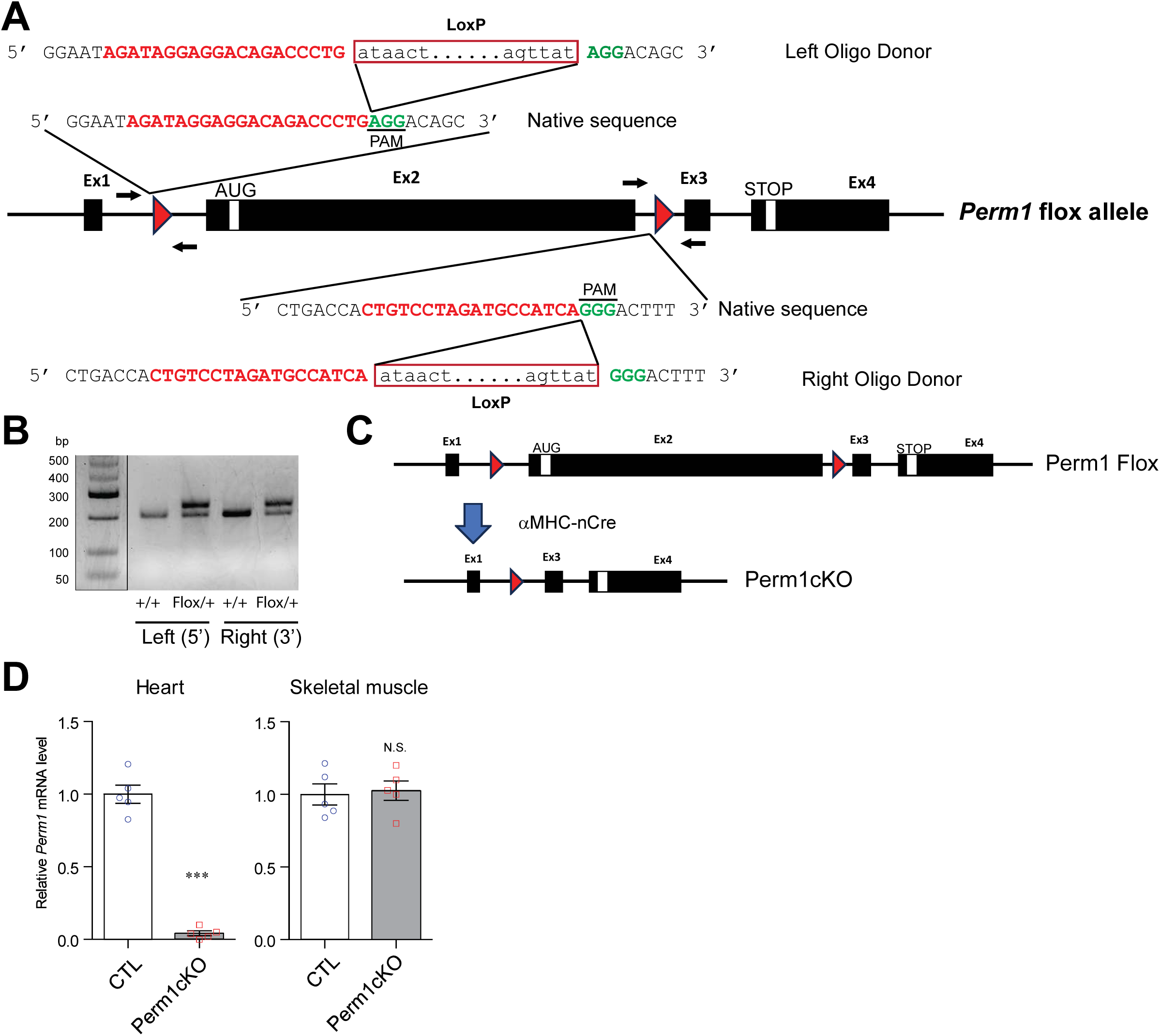
Generation of the Perm1 floxed allele and cardiomyocyte-specific Perm1 knockout mice. **(A)** Schematic of CRISPR/Cas9-mediated insertion of loxP sites flanking exon 2 of Perm1 to generate the conditional (floxed) allele, with representative PCR genotyping showing the size distinction between wild-type and floxed alleles. **(B)** PCR genotyping of tail DNA from Perm1^flox/flox mice and wild-type littermates. **(C)** Strategy for Cre-dependent deletion of exon 2 in cardiomyocytes using αMHC-nCre and the expected recombined allele. **(D)** RT-qPCR analysis of *Perm1* mRNA levels in heart (left ventricle) and skeletal muscle (gastrocnemius) from 2-month-old male control and cKO mice, confirming cardiac-specific deletion. Data are mean ± SEM; n = 5 per group. ***P < 0.001. Statistical analysis was performed using an unpaired Student’s t-test.

First, to generate a cardiomyocyte-specific Perm1 knockout (KO) model, Perm1^flox/flox mice were crossed with the cardiomyocyte-specific Cre driver αMHC-nCre [19] producing Perm1cKO mice (Perm1^flox/flox::αMHC-nCre) in which exon 2 is deleted in a Cre-dependent manner specifically in cardiomyocytes (**Fig. 1C**). To confirm tissue-restricted recombination, Perm1 mRNA levels were quantified by qPCR in heart and skeletal muscle (gastrocnemius) from 2-month-old male cKO mice and littermate controls. Perm1 mRNA was reduced by approximately 95% in cKO hearts compared with controls, whereas expression in skeletal muscle was unchanged, confirming cardiac-specific deletion (**Fig. 1D**).

### Perm1 is dispensable for cardiac development

We previously showed that Perm1 expression increases during cardiac development from the embryonic stage to adulthood, suggesting that Perm1 may contribute to cardiac maturation [12]. To determine whether cardiomyocyte-specific Perm1 deletion affects embryonic cardiac development, we first confirmed efficient deletion at the protein level. Immunoblotting of whole-heart lysates from embryonic day (E) 17.5 hearts demonstrated effective loss of Perm1 protein in cKO embryonic hearts (**Fig. 2A**).

**Figure 2.**
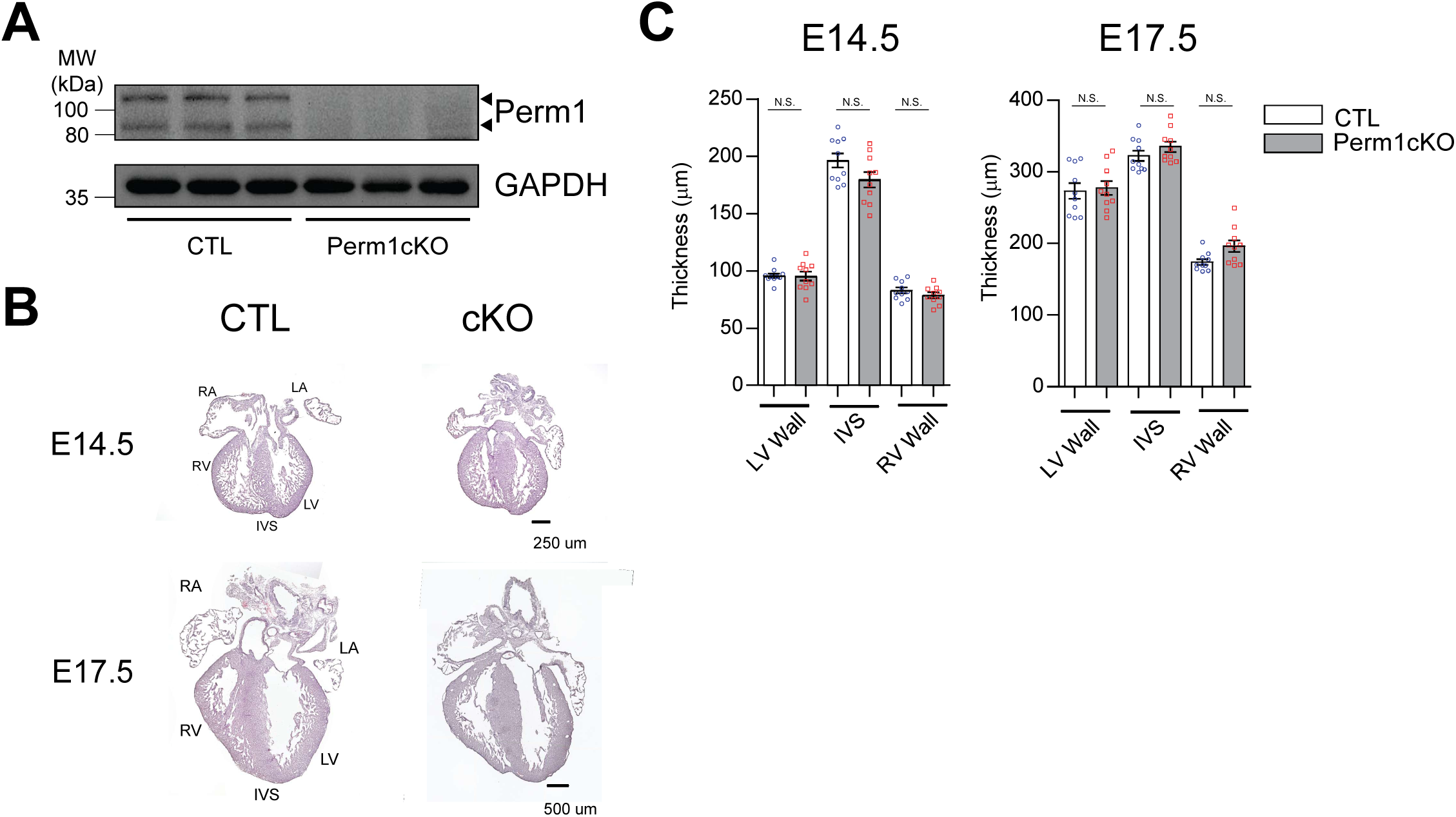
Cardiomyocyte-specific deletion of Perm1 does not affect embryonic cardiac development. Hearts from control and Perm1cKO embryos were analyzed at E14.5, when ventricular septation is essentially complete, and at E17.5, when ventricular compaction is well advanced. (A) Representative immunoblot of Perm1 in E17.5 whole-heart lysates, with GAPDH as a loading control. (B) Representative H&E-stained sections showing preserved ventricular architecture and chamber formation in both genotypes. (C) Quantification of left ventricular (LV) wall, right ventricular (RV) wall, and interventricular septum (IVS) thickness at E14.5 and E17.5. Data are mean ± SEM; Scale bars, 250 µm (E14.5) and 500 µm (E17.5). Statistical analysis was performed using an unpaired Student’s t-test.

We next analyzed cardiac morphology in control and Perm1cKO embryos at E14.5 and E17.5, corresponding to developmental stages at which ventricular septation is essentially complete and ventricular compaction is well advanced, respectively. Histological analysis by H&E staining showed preserved ventricular architecture and normal chamber formation at both time points (**Fig. 2B**). Quantification of left ventricular wall thickness, right ventricular wall thickness, and interventricular septal thickness revealed no significant differences between genotypes at either developmental stage (**Fig. 2C**).

### Deletion of Perm1 in the heart does not affect basal cardiac function

To assess whether cardiomyocyte-specific deletion of Perm1 alters basal cardiac function over time, we performed longitudinal cardiac phenotyping of Perm1cKO mice and littermate controls. No gross abnormalities of the mice were observed (**Fig. 3A**). Cardiac function was assessed by transthoracic echocardiography. Males were evaluated at 4, 8, and 12 months of age, whereas females were assessed at 12 months (**Fig. 3B-E, male; and Supplemental Fig. S1, female**). Global systolic performance, assessed by ejection fraction (EF) and fractional shortening (FS), as well as wall thickness and chamber dimensions, was preserved in cKO mice compared to controls at all time points examined. Pulse wave and tissue Doppler echocardiography [20] performed in 12-month-old male mice showed no significant differences in E/E′ or E/A ratios between control and cKO mice (**Supplemental Fig. S2**), indicating that diastolic function is also preserved.

**Figure 3.**
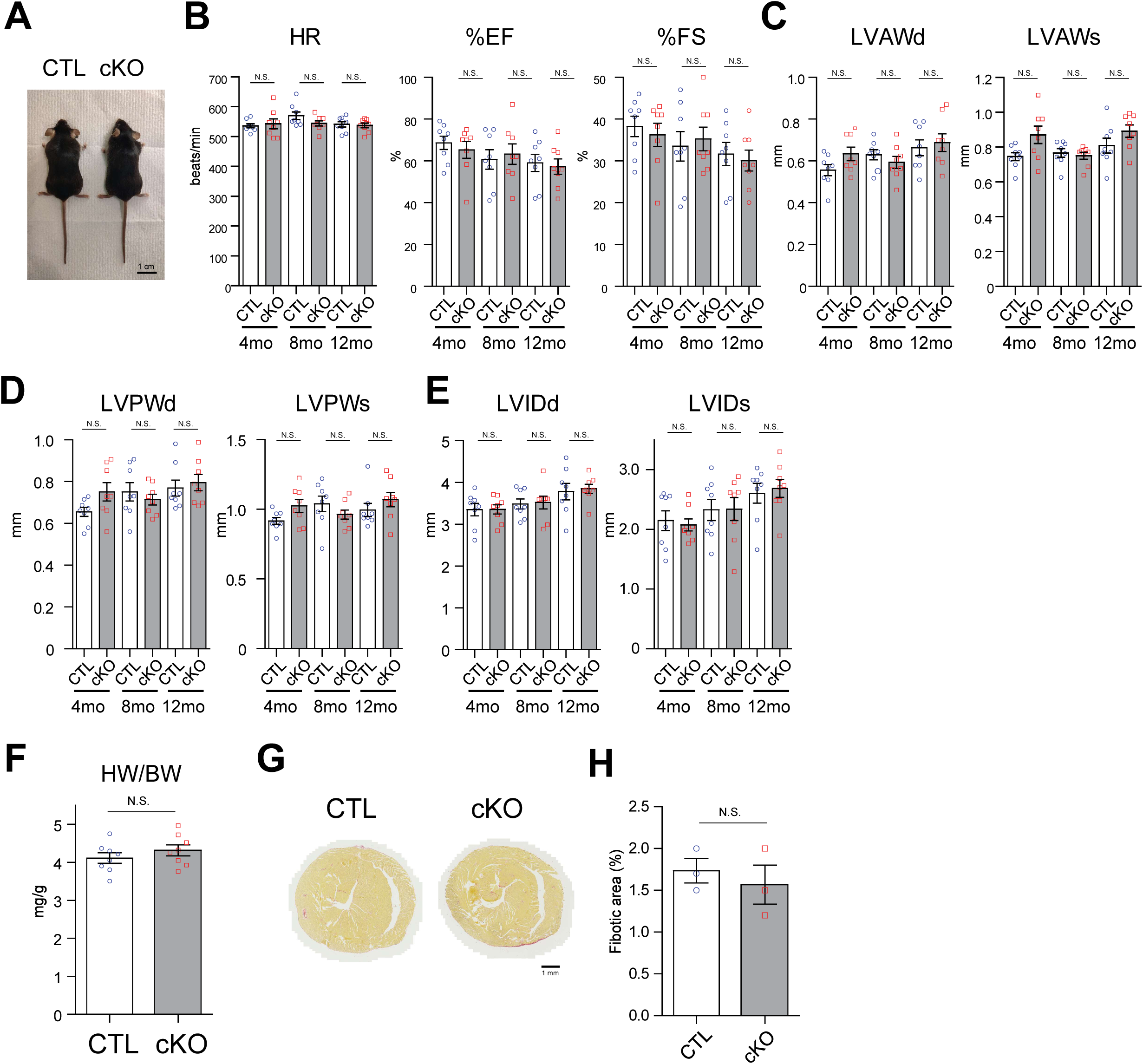
Constitutive cardiomyocyte-specific Perm1 deletion preserves basal cardiac structure and function through 12 months of age. Cardiac phenotype was evaluated in control (CTL) and Perm1cKO mice by transthoracic echocardiography. The male mice were assessed longitudinally at 4, 8, and 12 months of age; female mice were assessed at 12 months of age (Supplemental Fig. S1). **(A)** Representative images of 4-month-old male CTL and Perm1cKO mice. **(B)** Heart rate (HR), ejection fraction (EF), and fractional shortening (FS) in male mice at 4, 8, and 12 months of age. **(C)** Left ventricular anterior wall thickness in diastole (LVAWd) and systole (LVAWs). **(D)** Left ventricular posterior wall thickness in diastole (LVPWd) and systole (LVPWs). **(E)** Left ventricular internal dimension in diastole (LVIDd) and systole (LVIDs). **(F)** Heart weight-to-body weight ratio (HW/BW) in male mice at 12 months of age. **(G)** Representative Picrosirius Red-stained transverse sections of 12-month-old male hearts. **(H)** Quantification of fibrotic area, expressed as the percentage of Picrosirius Red-positive area relative to total myocardial area. Data are mean ± SEM. **(B-E)** n = 8-9 per genotype; **(F)** n = 8-9 per genotype; **(H)** n = 3 per genotype. Scale bars: 1 cm **(A)** and 1 mm **(G)**. Statistical analysis was performed using two-way ANOVA with genotype and time as factors, followed by Bonferroni’s multiple comparisons test **(B-E)**, or unpaired two-tailed Student’s *t* test **(F, H)**. N.S., not significant.

In addition to echocardiographic measurements, we observed no evidence of cardiac hypertrophy in cKO mice, as determined by normalized heart weight at 12 months (Fig. 3F). Picrosirius Red staining also showed no significant difference in fibrosis between control and cKO hearts at 12 months (**Fig. 3G and H**). Together, these results demonstrate that Perm1 in cardiomyocytes is dispensable for long-term maintenance of cardiac structure and function under basal conditions.

### Deletion of cardiomyocyte Perm1 in the adult heart does not affect basal cardiac function

Since the αMHC-nCre expresses constitutively in cardiac myocytes by mid-embryonic development [21] we generated an adult-inducible knockout model using the tamoxifen (TAMO)-inducible αMHC-MerCreMer driver [22]. Adult male mice (10-week-old) received tamoxifen (50 mg/kg body weight, i.p.) once daily for 5 consecutive days and were evaluated by echocardiography at baseline (pre-TAMO treatment) and at 2 and 4 months after the final injection; TAMO-treated Cre littermates served as experimental controls (**Fig. 4A**). The use of Cre-negative, tamoxifen-treated littermates as controls accounts for any potential effects of tamoxifen itself on cardiac function; we have previously found no evidence of tamoxifen- and MerCreMer-induced toxicity at this TAMO dosage [22].

**Figure 4.**
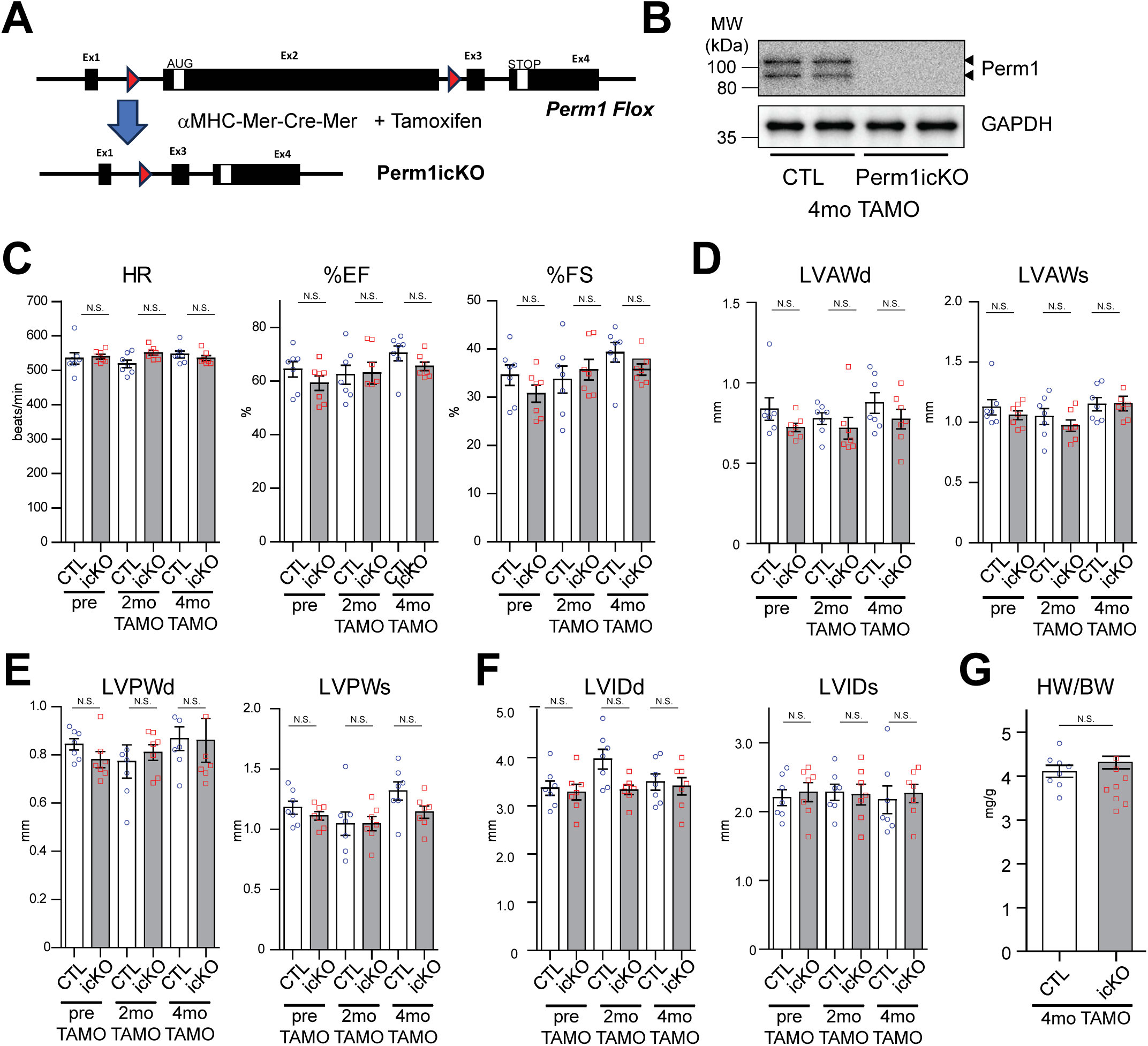
Adult-inducible deletion of Perm1 does not impair basal cardiac function. Perm1 was deleted in adult cardiomyocytes using the tamoxifen-inducible αMHC-MerCreMer driver. Ten-week-old male control (Perm1^flox/flox; Cre) and Perm1icKO (Perm1^flox/flox::αMHC-MerCreMer) mice received tamoxifen. (A) Schematic of Perm1icKO mouse generation. (**B**) Representative immunoblot of Perm1 and GAPDH (loading control) in adult LV lysates at 4 months post-injection. (**C**) Echocardiographic assessment of systolic function (HR, EF, and FS) at baseline and at 2 and 4 months post-injection. (**D-F**) Echocardiographic structural parameters (LVAWd, LVAWs, LVPWd, LVPWs, LVIDd, LVIDs). (**G**) Heart weight-to-body weight ratio (HW/BW) at 4 months post-injection. Data are mean ± SEM; n = 7-8 per group. Statistical analysis used two-way ANOVA with genotype and time as factors, followed by Bonferroni’s multiple comparisons test (**C-F**), or an unpaired two-tailed Student’s *t* test **(G)**. N.S., not significant.

Immunoblotting confirmed efficient loss of Perm1 protein in adult Perm1icKO hearts at 4 months after TAMO injection (**Fig. 4B**). Despite robust Perm1 deletion, systolic function remained preserved, with EF and FS comparable between icKO mice and controls at both post-induction time points (**Fig. 4C**). Cardiac dimensions and wall structure parameters were also unchanged (**Fig. 4D-F**), and normalized heart weight at 4 months post-induction did not differ between genotypes (**Fig. 4G**). Together, consistent with the constitutive αMHC-nCre-driven Perm1cKO model, these data show that adult-onset Perm1 deletion does not impair basal cardiac function.

### Perm1 deletion reduces mitochondrial respiration and alters mitochondrial organization without changes in total mitochondrial abundance

We previously reported that Perm1 augmentation promotes mitochondrial biogenesis and oxidative metabolism in cardiomyocytes [12]. Because basal cardiac structure and function were preserved in both sexes (**Fig. 3** and **Supplemental Fig. S1**), subsequent mechanistic analyses were performed in male mice. To determine whether Perm1 deletion alters cardiac mitochondrial bioenergetics, we assessed mitochondrial respiration in heart tissue from 6-month-old control and Perm1cKO mice using high-resolution respirometry (Oroboros O2k) (**Fig. 5A**). Respiration was first recorded in the absence of exogenous substrates. We then measured respiration sequentially with glutamate/malate (GM), followed by ADP to stimulate oxidative phosphorylation, and succinate to engage combined complex I+II-supported respiration. Under these conditions, mitochondrial respiration in cKO hearts was reduced by approximately 20-25% compared with controls: GM-supported respiration was decreased by ∼25% (p < 0.05), complex I-driven OxPhos (GM+ADP) by ∼20% (p < 0.05), and combined complex I+II-supported OxPhos (GM+ADP+succinate) by ∼25% (p < 0.05) (**Fig. 5A**). The observation that succinate-driven complex II input did not restore respiration to control levels suggests that the respiratory deficit is not restricted to complex I but reflects a broader impairment in mitochondrial oxidative capacity.

**Figure 5.**
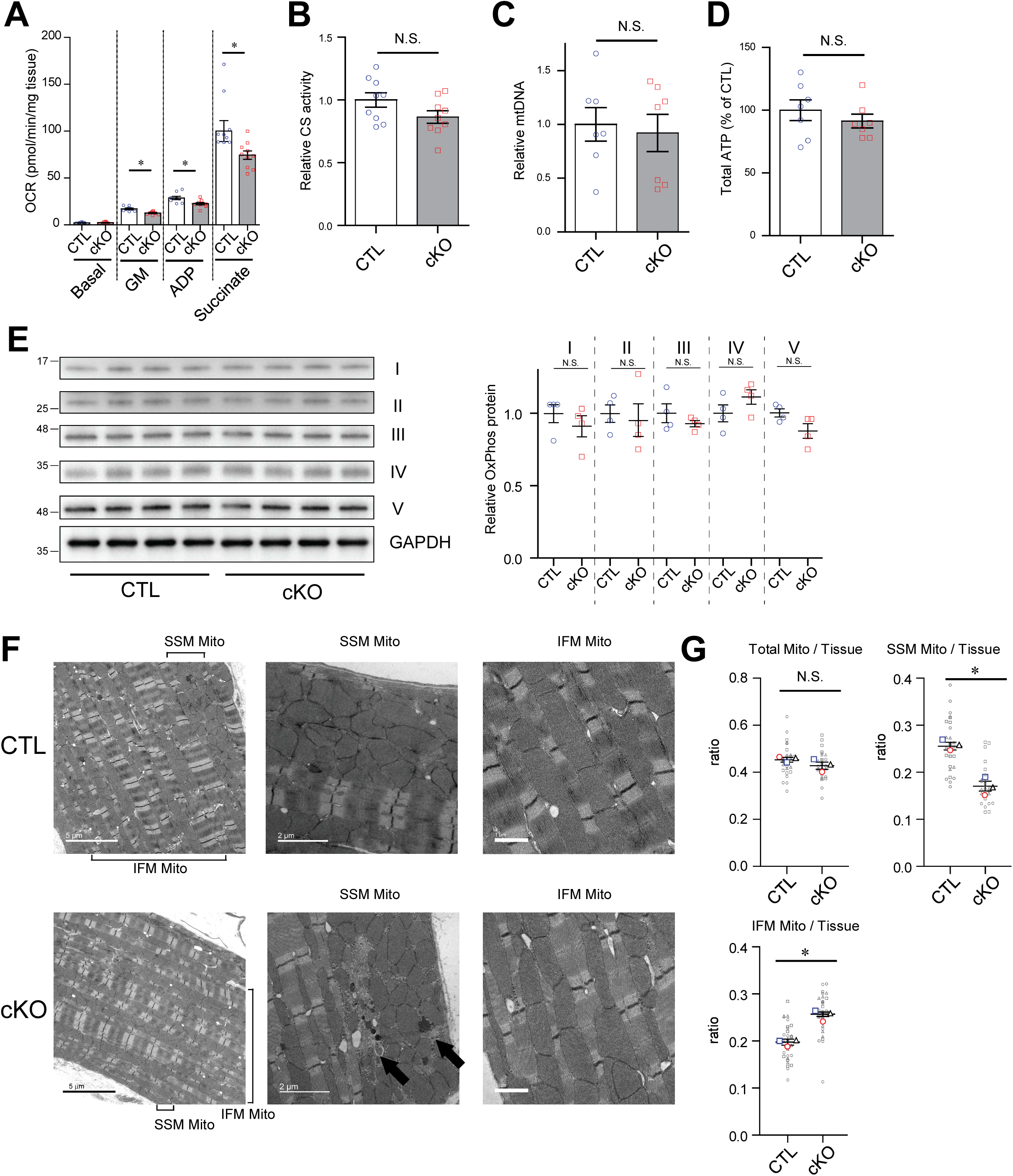
Perm1 deletion reduces cardiac mitochondrial respiration and alters mitochondrial distribution without changing total mitochondrial content. LV tissues were harvested from 6-month-old male control (CTL) and Perm1cKO mice. (A) Oxygen consumption rate (OCR) assessed in permeabilized LV tissue using high-resolution respirometry (Oroboros O2k). Measurements were recorded sequentially under the following conditions: no exogenous substrate (Basal), complex I substrates glutamate/malate (GM), GM + ADP (complex I-driven OxPhos), and GM + ADP + succinate (combined complex I+II-supported OxPhos). OCR was normalized to tissue weight (mg). (B) Citrate synthase (CS) activity, expressed relative to CTL. (C) Relative mitochondrial DNA content, calculated as the ratio of mitochondrial (CoxII) to genomic (Dio3) DNA copy number and expressed relative to CTL. (D) Total ATP content, normalized to tissue protein and expressed relative to CTL. (E) Immunoblot analysis of representative OxPhos subunits (complexes I-V) in LV lysates, with GAPDH as a loading control. Quantification is shown on the right, with CTL set to 1. (F) Representative transmission electron microscopy (TEM) images of LV myocardium. Upper panels, CTL; lower panels, cKO. Left, subsarcolemmal (SSM) region at low magnification; center, SSM region at high magnification; right, interfibrillar (IFM) region. Arrows indicate structures morphologically consistent with mitochondria-containing autophagosomes in the subsarcolemmal region of cKO hearts. Scale bars, 5 µm (left), 2 µm (center), and 1 µm (right). (G) Quantification of mitochondrial area relative to tissue area, including total mitochondria/tissue, subsarcolemmal mitochondria (SSM) / tissue, and interfibrillar mitochondria (IFM) / tissue ratios. Small open symbols denote individual fields and large filled symbols denote the mean of each mouse, with symbol shape identifying the animal; bars show mean ± SEM of the animal means. (A) n = 9-10; (B) n = 8-9; (C) n = 7; (D) n = 7; (E) n = 4 per genotype; (F, G) n = 3 mice per genotype, with 10-12 non-overlapping fields analyzed per group. Statistical analysis was performed using an unpaired Student’s t-test; for (G), statistics were performed on the animal means. *P < 0.05. N.S., not significant.

To determine whether reduced respiration reflected decreased mitochondrial content, we quantified citrate synthase activity, mitochondrial DNA copy number, and total ATP content. Citrate synthase activity and mtDNA copy number were comparable between control and cKO hearts (**Fig. 5B-C**), and total ATP levels were also unchanged (**Fig. 5D**). We further examined OxPhos protein levels by immunoblotting; the abundance of representative OxPhos subunits (complexes I-V) was similar between genotypes (**Fig. 5E**). Together, these findings indicate that the observed respiratory deficit in cKO hearts is not attributable to reduced mitochondrial mass or global loss of respiratory chain components.

Despite preserved total mitochondrial content, electron microscopy revealed remodeling of mitochondrial subpopulations. Quantification of the mitochondria-to-tissue area ratio showed no difference in total mitochondrial density between genotypes (**Fig. 5G**). However, the subsarcolemmal mitochondrial fraction was significantly reduced (∼30%) in cKO hearts, whereas interfibrillar mitochondria were significantly increased (∼30%) (**Fig. 5F-G**), indicating a redistribution of mitochondrial subpopulations rather than a net gain or loss. In addition, occasional structures morphologically consistent with mitochondria-containing autophagosomes were noted in the subsarcolemmal region of cKO hearts (**Fig. 5F**).

Together, these data demonstrate that cardiomyocyte-specific Perm1 deletion reduces mitochondrial respiratory capacity by approximately 20-25% and is associated with spatial redistribution of mitochondrial subpopulations, without detectable changes in total mitochondrial abundance or OxPhos protein content.

### Molecular changes associated with Perm1 deletion in the heart

We previously reported that Perm1 enhances PGC-1/ERR pathway activity and MIC60/IMMT protein expression, promoting oxidative metabolism in cardiomyocytes and the heart [12; 13]. To investigate molecular changes associated with Perm1 deletion, we examined the expression of key transcriptional regulators, metabolic enzymes, and hypertrophic markers in 6-month-old male Perm1cKO hearts.

qPCR analysis revealed selective changes in the PGC-1/ERR transcriptional network. Perm1 mRNA was reduced by approximately 95% (p < 0.001), confirming efficient deletion (**Fig. 6A**). Among transcriptional regulators, PGC-1α was significantly reduced (p < 0.05), as were ERRβ and PPARα (p < 0.05 for both), whereas ERRγ was notably increased (p < 0.05) (**Fig. 6A**). The upregulation of ERRγ in the context of reduced PGC-1α and ERRβ may reflect activation of a compensatory transcriptional program aimed at partially sustaining oxidative gene expression in the absence of Perm1. Expression of other transcriptional regulators and mitochondrial biogenesis factors, including PGC-1β, ERRα, PPARδ, TFAM, NRF-1, NRF-2, and Mfn2, was unchanged between genotypes (**Fig. 6A-B**). Similarly, glucose metabolism genes (GLUT4, LDHb, PDK4), fatty acid metabolism genes (Lpl, FABP3, CPT1b, LCAD, ATGL), and mitochondrial enzyme and electron transport chain genes (Ckmt2, IDH3a, COX7a1) did not differ significantly (**Fig. 6C-E**). Hypertrophic markers ANF and BNP, as sensitive markers of myocardial stress, were also unchanged, consistent with the absence of echocardiographic changes and cardiac hypertrophy in cKO mice (**Fig. 6F**).

**Figure 6.**
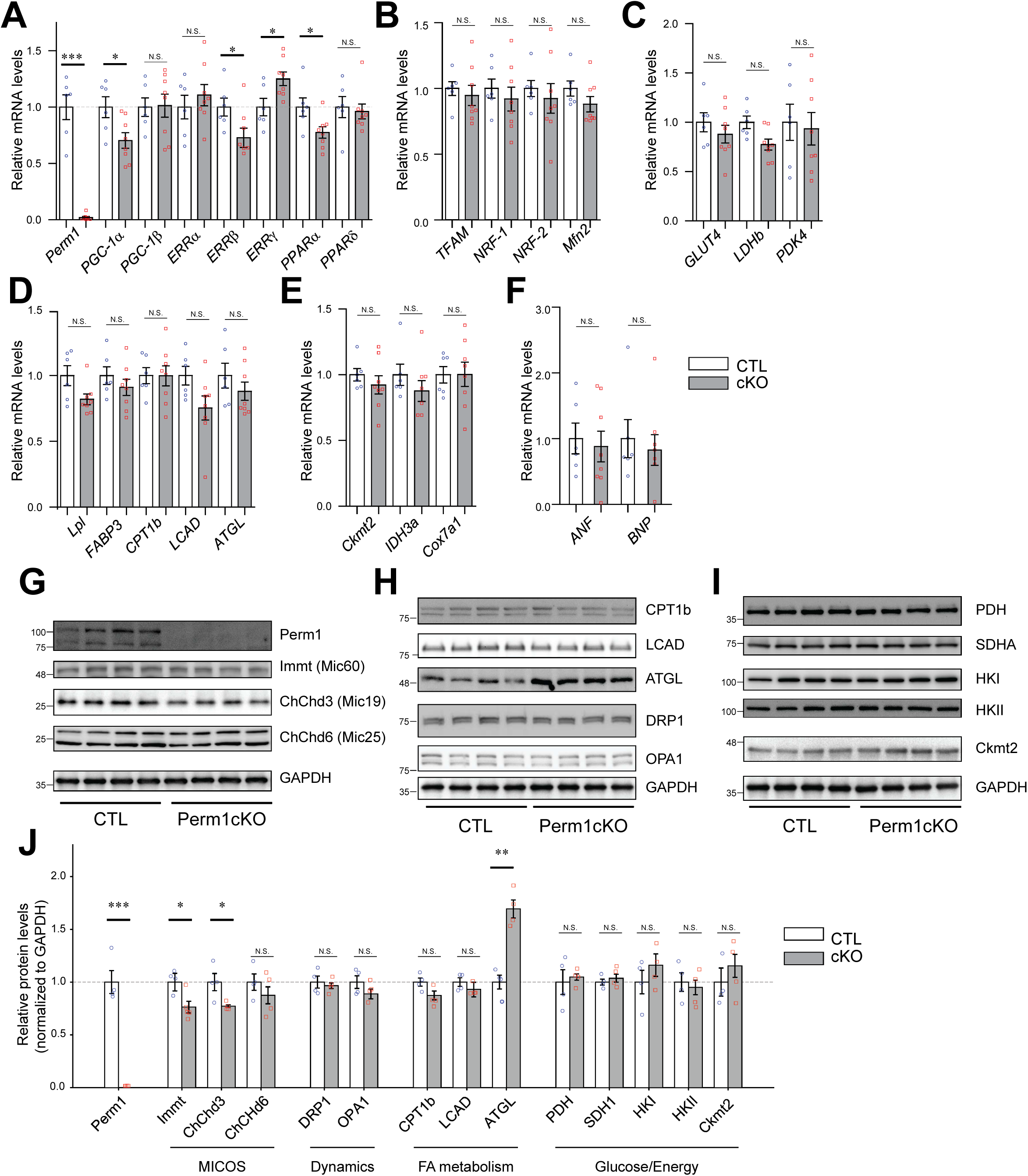
Perm1 deletion induces selective transcriptional and protein-level remodeling of metabolic and mitochondrial programs. LV tissues from 6-month-old male control and cKO mice were analyzed for gene expression and protein abundance. (**A**) qPCR analysis of transcriptional regulators of oxidative metabolism, including Perm1, PGC-1α/β, ERRα/β/γ, and PPARα/δ. (**B**) Mitochondrial biogenesis factors (TFAM, NRF-1, NRF-2, Mfn2). (**C**) Glucose metabolism genes (GLUT4, LDHb, PDK4). (**D**) Fatty acid metabolism genes (Lpl, FABP3, CPT1b, LCAD, ATGL). (**E**) Mitochondrial enzyme and electron transport chain genes (Ckmt2, IDH3a, COX7a1). (**F**) Hypertrophic markers (ANF, BNP). All mRNA levels are expressed relative to CTL, with CTL set to 1. (**G**) Immunoblot analysis of Perm1 and MICOS proteins (Perm1, Immt(Mic60), ChChd3(Mic19), ChChd6(Mic25)) with GAPDH as a loading control. (**H**) Immunoblot analysis of lipid metabolic enzymes and mitochondrial dynamics proteins (CPT1b, LCAD, ATGL, DRP1, OPA1) with GAPDH as a loading control. (**I**) Immunoblot analysis of metabolic enzymes involved in energy and glucose metabolism (PDH, SDHA, HKI, HKII, Ckmt2) with GAPDH as a loading control. (**J**) Quantification of protein abundance from (G-I). Protein levels are expressed relative to CTL, with CTL set to 1. Data are mean ± SEM; n = 8-9 (A-F) or 4 (**G-J**) per group. *P < 0.05, **P< 0.01, ***P < 0.001. Statistical analysis was performed using an unpaired Student’s t-test.

At the protein level, immunoblotting revealed that Perm1 protein was markedly reduced in cKO hearts, confirming efficient deletion (**Fig. 6G, J**). Among the proteins assessed, we identified three significant changes. First, IMMT (MIC60), a core component of the MICOS complex that regulates cristae junction formation[23], was modestly but significantly reduced (p < 0.05) (**Fig. 6G, J**). Second, ChChd3 (MIC19), another MICOS-associated protein [24], was also significantly reduced (p < 0.05) (**Fig. 6G, J**). Third, ATGL protein abundance was significantly increased (p < 0.01) (**Fig. 6H, J**), despite unchanged ATGL mRNA levels (**Fig. 6D**), suggesting post-transcriptional or post-translational regulation of ATGL in Perm1-deficient hearts. The remaining targets, including DRP1, OPA1, ChChd6 (MIC25), PDH, SDHA, Ckmt2, HKI, HKII, CPT1b, and LCAD, were comparable between genotypes (**Fig. 6G-J**).

Together, these data reveal a selective molecular signature in Perm1-deficient hearts: partial downregulation of the PGC-1α/ERRβ/PPARα transcriptional axis, concurrent increase in ERRγ, reduced abundance of MICOS complex components IMMT and ChChd3, and elevated ATGL protein.

### Perm1cKO mice show reduced exercise capacity

During exercise, the heart must rapidly increase ATP production to sustain augmented contractile performance and systemic oxygen delivery. Given the mitochondrial respiratory deficit and ultrastructural remodeling observed in Perm1-deficient hearts, we tested whether Perm1 deletion in cardiac myocytes limits exercise capacity. We performed treadmill running-to-exhaustion tests in male and female mice at 6 and 12 months of age using age-appropriate protocols: a graded protocol with stepwise speed increments (5, 10, 15, and 20 m/min) at 6 months, and a simplified two-stage protocol (5 and 10 m/min) at 12 months to accommodate age-related differences in exercise capacity (**Fig. 7A**). The 6 month old cKO mice exhibited significantly reduced exercise performance compared with controls, with decreased total running time and distance to exhaustion in both males and females (**Fig. 7B and C**). This impairment persisted at 12 months of age, as cKO mice continued to show significantly lower running time and distance than controls in both sexes (**Fig. 7B and C**).

**Figure 7.**
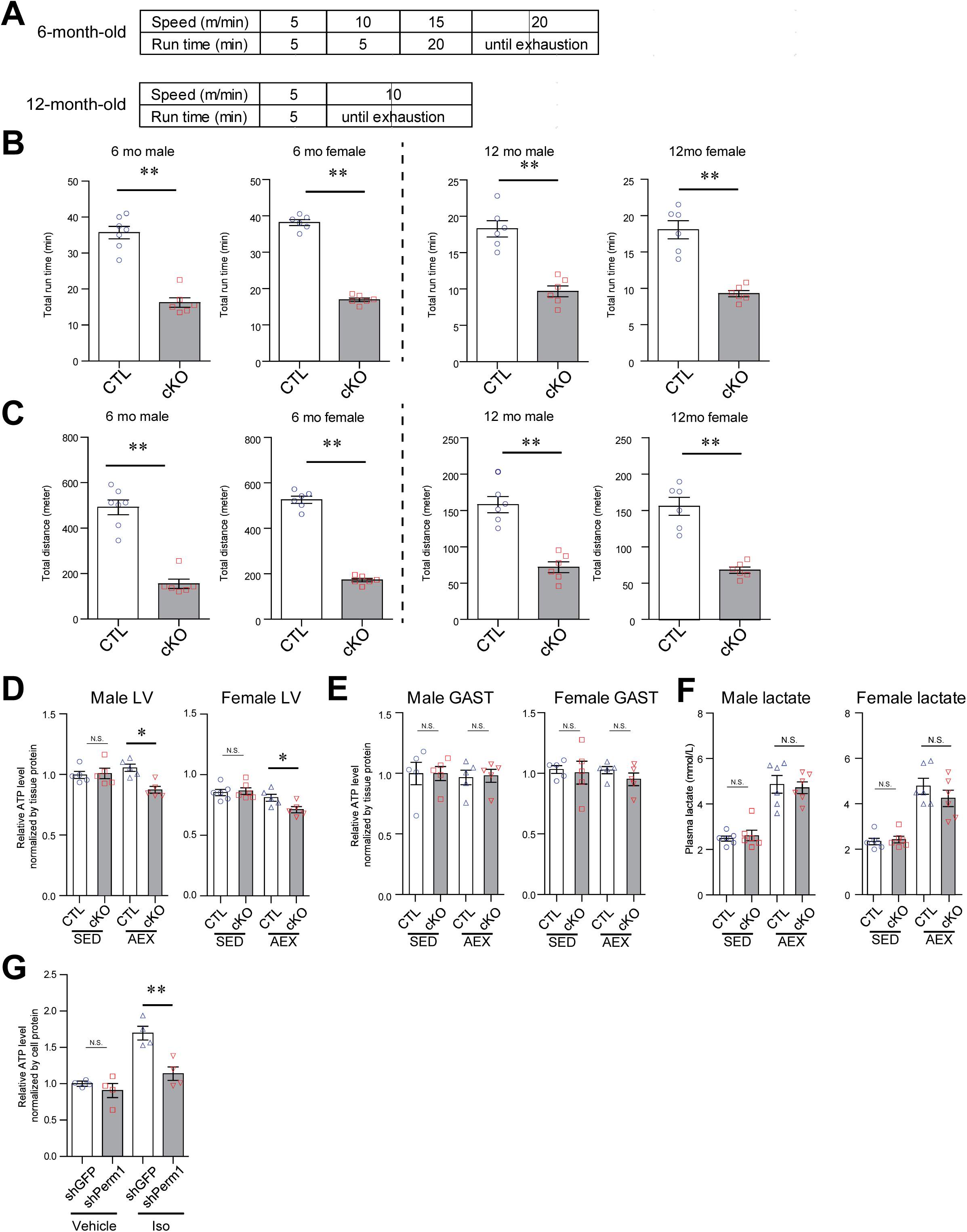
Cardiac-specific deletion of Perm1 impairs exercise capacity and post-exercise cardiac ATP maintenance. (**A**) Treadmill protocols for running-to-exhaustion tests in male and female control (CTL) and Perm1cKO mice at 6 and 12 months of age. At 6 months, treadmill speed increased stepwise (5, 10, 15, and 20 m/min); at 12 months, a simplified two-stage protocol (5 and 10 m/min) accommodated age-related differences in baseline running capacity. (**B**) Total running time (left) to exhaustion in male and female mice at 6 (left) and 12 (right) months of age. (C) Total distance run to exhaustion in male and female mice at 6 (left) and 12 (right) months of age. (**D**) Total ATP content in left ventricular tissue from 6-month-old male and female mice under SED (sedentary) and AEX (post-exercise) conditions, normalized to tissue protein and expressed relative to SED CTL. (**E**) Total ATP content in gastrocnemius muscle from 6-month-old male and female mice under SED and AEX conditions, normalized to tissue protein and expressed relative to SED CTL. (**F**) Plasma lactate levels in 6-month-old male and female mice under sedentary (SED) and post-exercise (AEX) conditions. (**G**) Relative ATP content in cultured neonatal mouse cardiomyocytes transduced with shRNA targeting Perm1 (shPerm1) or GFP (shGFP; control), treated with vehicle or isoproterenol (Iso; 1 µM) for 30 min. ATP content was normalized to total cellular protein and expressed relative to vehicle-treated shGFP cells. Data are mean ± SEM. (B-F) n = 5-6 per sex/genotype/age; (G) n = 4 per group. Statistical analysis was performed using unpaired two-tailed Student’s *t-test* (B and C) or two-way ANOVA followed by Bonferroni’s multiple comparisons test (D-G). \**P* < 0.05; \*\**P* < 0.01; N.S., not significant.

To determine whether the exercise phenotype was associated with cardiac bioenergetic limitation, we quantified total ATP content in hearts and gastrocnemius (GAST) muscles from 6-month-old mice under both sedentary (SED) and post-exercise (AEX) conditions. For post-exercise sampling, mice underwent a 12-min running protocol (5 m/min for 5 min, 10 m/min for 5 min, then 15 m/min for 2 min), a time point selected to precede the onset of exhaustion in cKO mice, enabling comparison of cardiac energetics at a matched submaximal workload rather than at different stages of fatigue. Under sedentary conditions, cardiac ATP levels were comparable between control and cKO hearts (**Fig. 7D**), consistent with preserved basal cardiac function. Following exercise, however, cardiac ATP levels were significantly reduced in cKO hearts compared with controls in both sexes (**Fig. 7D**). In contrast, ATP levels in GAST muscle were unchanged between genotypes under both sedentary and post-exercise conditions (**Fig. 7E**), indicating that the bioenergetic deficit is cardiac-specific rather than reflecting a skeletal muscle impairment. Plasma lactate levels measured after the 12-min run were also comparable between control and cKO mice (**Fig. 7F**), suggesting that the reduced running capacity in cKO mice is unlikely to reflect differences in peripheral anaerobic metabolism.

To determine whether this energetic limitation is cardiomyocyte-autonomous, we examined the acute metabolic response to β-adrenergic stimulation, the principal signal coupling exercise to increased cardiac workload, in cultured neonatal mouse cardiomyocytes. Cells were transduced with shRNA targeting Perm1 (shPerm1) or GFP (shGFP) as a control and treated with isoproterenol (1 µM, 30 min), a maximal β-adrenergic challenge used as an in vitro surrogate for the catecholamine-driven increase in cardiac workload during exercise [25] [26]. Knockdown efficiency was confirmed in parallel cultures, with Perm1 reduced by approximately 80% in shPerm1-transduced cardiomyocytes relative to shGFP controls (Supplemental Fig. S3).

Isoproterenol selectively engages β-adrenergic signaling without concurrent activation of α1-adrenergic receptors, which are also expressed in neonatal cardiomyocytes and would confound interpretation of β-adrenergic-driven metabolic responses. The isoproterenol concentration was selected based on preliminary dose-optimization experiments (1, 10, and 100 µM), in which 1 µM produced the maximal increase in cellular ATP content, consistent with β-adrenergic receptor desensitization at supramaximal agonist concentrations [27].

Under basal conditions, ATP content did not differ between shGFP and shPerm1 cardiomyocytes (P > 0.99). Isoproterenol significantly increased ATP content 1.70-fold in shGFP cells (to 1.70 ± 0.09, P < 0.001), whereas this response was blunted in shPerm1 cells (1.25-fold, to 1.14 ± 0.09, P = 0.29 vs its own vehicle control), resulting in significantly lower ATP levels in Perm1-knockdown cardiomyocytes under stimulated conditions (P < 0.01) (**Fig. 7G**). Two-way ANOVA revealed a significant genotype × isoproterenol interaction (F(1,12) = 8.04, P = 0.015), indicating that Perm1 knockdown selectively impairs the capacity to augment ATP production in response to adrenergic stimulation.

Together, these data demonstrate that cardiomyocyte-specific Perm1 deletion compromises exercise tolerance in both sexes across multiple ages.

## 3. Discussion

In this study, we used two complementary cardiomyocyte-specific knockout strategies to define Perm1’s cell-autonomous function in the heart. Constitutive deletion via αMHC-nCre established that Perm1 is dispensable for embryonic cardiac development and for maintaining cardiac structure and function through 12 months of age. An adult tamoxifen-inducible model confirmed that this preserved phenotype does not reflect developmental compensation. Despite normal basal parameters and resting performance, Perm1-deficient mice of both sexes showed reduced exercise capacity at multiple ages and failed to maintain cardiac ATP during exercise. Likewise, Perm1-deficient neonatal myocytes failed to augment ATP production upon β-adrenergic stimulation. Perm1 is therefore a cardiomyocyte-intrinsic regulator of bioenergetic reserve that becomes essential when energy demand exceeds resting levels.

Our data indicate that Perm1 contributes to the subsarcolemmal positioning of mitochondria in the heart, as it does in skeletal muscle. Perm1 is proposed to be an important bridge between mitochondria and their subcellular positioning in skeletal muscle [28]. Specifically, in skeletal muscle, PERM1 physically associates with the MICOS-MIB complex, which allows physical connection of mitochondria to the sarcolemma via ankyrin B [28], and mice with KO of Perm1 in skeletal muscle demonstrate reduced mitochondrial membrane potential, reduced complex I activity, and fewer SSM [28]. Our data extend these observations to cardiac muscle, as we find a 30% reduction in SSM in the LV from Perm1cKO mice, accompanied by a decrease in the MICOS components IMMT (MIC60) and ChChd3 (MIC19). Thus, these data suggest that Perm1 is an important (albeit partial) contributor to mitochondrial localization to the subsarcolemmal region.

This redistribution does not, however, account for the reduced respiration we measured. Because IFM are more respiratory-competent than SSM [29], a shift toward IFM would be expected to preserve, rather than reduce, tissue respiratory capacity [30]. Citrate synthase activity, mtDNA copy number, and OxPhos subunit abundance were likewise unchanged, so the 20-25% deficit cannot be attributed to a loss of mitochondrial mass or of respiratory chain components. The concurrent depletion of IMMT (MIC60) and ChChd3 (MIC19), which together form the core MICOS subcomplex that stabilizes crista junctions [24], points instead to altered inner membrane organization. Loss of MICOS integrity redistributes respiratory complexes within the inner membrane without preventing their assembly [23], which would reduce respiratory capacity while leaving subunit abundance intact - the pattern we observe. In parallel, partial downregulation of PGC-1α, ERRβ, and PPARα was accompanied by increased ERRγ transcript and ATGL protein, without any change in ATGL mRNA, suggesting compensatory transcriptional and lipid-mobilizing programs [31]. Together, these changes describe a state in which compensation suffices to maintain basal energetic balance but fails when ATP demand rises acutely.

The exercise phenotype observed in vivo in our mouse model could arise from cardiac limitation, skeletal muscle dysfunction, or systemic factors [32] [33] [34]. Three observations argue for a cardiac origin. Post-exercise ATP depletion was restricted to the heart, with gastrocnemius ATP unchanged; plasma lactate was comparable between genotypes, arguing against a difference in peripheral anaerobic metabolism; and most directly, Perm1 knockdown in cultured neonatal cardiomyocytes recapitulated the deficit in a cell-autonomous setting, with β-adrenergic stimulation raising ATP content 1.7-fold in control cells but only 1.25-fold after Perm1 depletion, with no difference under basal conditions. Because β-adrenergic signaling is the principal stimulus coupling exercise to increased cardiac workload, this links the in vivo phenotype to a defined cell-autonomous mechanism.

This pattern of normal basal function with selectively impaired capacity to expand energy output parallels observations in brown adipocytes lacking all three estrogen-related receptors, which maintain normal basal respiration yet fail to increase spare respiratory capacity upon adrenergic stimulation [4]. Given that Perm1 is itself a PGC-1/ERR-responsive gene, our findings position Perm1 as a candidate effector through which this transcriptional axis confers adrenergic-responsive bioenergetic reserve in striated muscle. Notably, the ERR-dependent responses in brown adipocytes were transcriptional and required hours of stimulation [4], whereas the deficit we observe emerges within 30 minutes, indicating that Perm1 also supports acute metabolic responsiveness that transcriptional remodeling alone cannot explain.

In the healthy heart, the creatine kinase system holds ATP concentration essentially constant across workloads, and phosphocreatine declines before ATP does [6; 34]. Even in diseased human hearts, exercise typically unmasks an energetic deficit as a fall in PCr/ATP rather than in ATP itself [6; 35–37]. Total cardiac ATP fell in Perm1cKO hearts at a submaximal workload, at a time point preceding exhaustion, indicating a deficit that had already exceeded the phosphagen system’s buffering capacity.

This study did not directly determine how this deficit limits exercise capacity, but several non-exclusive consequences may apply. Cardiac ATP is consumed principally by myosin ATPase, SERCA2a, and Na /K -ATPase, and a fall in ATP would constrain both contractile and relaxation reserve [7; 38]. Relaxation may be the more vulnerable of the two: calcium reuptake by SERCA2a is directly ATP-dependent, and diastolic filling time is progressively curtailed as heart rate rises, so delayed relaxation would compromise ventricular filling precisely when stroke volume must increase [7]. Two features of our data support this possibility: diastolic parameters were normal at rest through 12 months, indicating a load-dependent rather than constitutive deficit; and SSM area was selectively decreased. SSM and IFM both supply membrane-associated ion transport, and SSM is more susceptible than IFM to calcium-mediated inhibition of ATP synthesis[29; 39–41]. Therefore, during exercise, the most vulnerable population faces the greatest demand. Because the heart extracts 70-80% of delivered oxygen even at rest [32], any constraint on myocardial ATP production translates into a constraint on cardiac output and hence on oxygen delivery to working muscle. We did not, however, measure cardiac output or contractile reserve during exercise, and this link therefore remains inferential.

Our findings place Perm1 within a broader pattern among components of the PGC-1/ERR axis. Combined or upstream loss produces severe phenotypes: cardiac deletion of both PGC-1α and PGC-1β causes perinatal heart failure with arrested mitochondrial biogenesis [5], and ERRγ-null mice fail to complete the postnatal switch to oxidative metabolism and die within the first week of life [42]. In contrast, loss of individual upstream regulators is largely compensated at rest but decompensates under load. ERRα-null mice are viable with no baseline cardiac phenotype, yet develop chamber dilatation and systolic dysfunction under pressure overload, with abnormal phosphocreatine depletion and reduced maximal ATP synthesis rates, a defect the authors described as impaired ATP reserve [43]. Perm1 deletion represents the most selective version of this phenotype: cardiac development, resting structure, and resting function are entirely preserved, and only the capacity to expand ATP production is lost. This gradient, from lethal disruption of mitochondrial biogenesis upstream to conditional loss of reserve capacity downstream, suggests that the PGC-1/ERR axis governs not a single output but a hierarchy, in which distal effectors such as Perm1 set the adaptive margin rather than the baseline. ERR-deficient brown adipocytes show an analogous dissociation, with normal basal respiration but absent spare respiratory capacity [4], indicating that this organization is not restricted to the heart.

Global and cardiomyocyte-specific deletion address different questions, and our findings are not incompatible with previous reports. Cardiomyocyte-specific deletion preserved basal cardiac function through 12 months in both constitutive and inducible models, with a sample size (n = 8-9) providing approximately 80% power to detect an effect of the magnitude previously reported for global deletion (Cohen’s d ≈ 1.4) [14]. Because global Perm1 loss also reduces skeletal muscle function [28], extracardiac contributions may add to the cardiac phenotype in whole-body models. More fundamentally, our data suggest that Perm1 loss compromises reserve capacity while basal function remains compensated. If so, whether resting dysfunction becomes detectable would depend on the cumulative demand a given cohort has experienced. This may explain why systolic impairment has been observed in some studies and not others, and would predict that the deficit becomes apparent as compensation is progressively exhausted.

Several limitations should be noted. Although our molecular and ultrastructural findings are consistent with a MICOS-dependent mechanism, they remain correlative; establishing causality will require direct morphometric assessment of crista architecture and tests of whether MIC60 restoration rescues the respiratory deficit [44]. Similarly, we did not examine whether the PERM1-ankyrin B interaction governs SSM positioning in cardiomyocytes as it does in skeletal muscle [28]. We observed occasional structures morphologically consistent with mitochondria-containing autophagosomes but did not assess mitophagy flux, leaving open whether SSM loss reflects enhanced turnover. Finally, our in vitro experiments used neonatal cardiomyocytes, which are more glycolytic than adult cells [45], and the isoproterenol concentration required to elicit a robust response exceeds plasma catecholamine levels attained during exercise[46]. This stimulation should therefore be regarded as a maximal β-adrenergic challenge rather than a mimic of exercise. Addressing these questions will require exercise stress echocardiography to determine whether contractile or relaxation function is impaired in vivo.

### Conclusions

These findings identify Perm1 as a cardiomyocyte-intrinsic determinant of cardiac bioenergetic reserve. Showing that resting cardiac function can be preserved while the capacity to expand ATP production is compromised has implications beyond Perm1 itself: conventional measures of contractile function may appear normal in hearts whose ability to respond to stress is already diminished. Identifying the effectors that confer this reserve, and determining whether they can be pharmacologically augmented, may offer an approach to cardiac metabolic disease that is not captured by resting phenotyping.

## 4. Materials and Methods

### 4.1 Animals

All animal studies were performed in accordance with the Guide for the Care and Use of Laboratory Animals (National Academies Press, 8th edition, 2011) and were approved by the Institutional Animal Care and Use Committee (IACUC) at the University of California, San Diego. All mice were maintained on a C57BL/6 background under a 12-h light/dark cycle at constant temperature (22 ± 2°C), with food and water provided ad libitum.

### -Generation of Perm1 floxed mice

LoxP sites were inserted to flank exon 2 of Perm1 using CRISPR/Cas9-mediated genome editing. Methods for CRISPR target sequence design, zygote microinjection, PCR screening, and sequence analysis have been described previously [18]. Correct targeting and cis integration of loxP sites were confirmed by PCR genotyping and genomic sequencing. All primers used in this study are listed in Supplemental Table 1.

### -Generation of cardiomyocyte-specific Perm1 knockout mice

Perm1^flox/flox mice were crossed with αMHC-nCre mice [19] to generate cardiomyocyte-specific Perm1 knockout mice (Perm1^flox/flox::αMHC-nCre; hereafter Perm1cKO). Littermate Perm1^flox/flox mice lacking Cre were used as controls unless otherwise indicated.

### -Tamoxifen-inducible Perm1 knockout mice

Perm1^flox/flox mice were crossed with αMHC-MerCreMer mice to generate Perm1^flox/flox::αMHC-MerCreMer mice (hereafter Perm1icKO). Ten-week-old male Perm1^flox/flox and Perm1icKO mice received tamoxifen (50 mg/kg body weight, i.p.) once daily for 5 consecutive days to induce Perm1 deletion in the adult heart [22]. Tamoxifen-treated Cre-negative littermates served as controls to account for potential effects of tamoxifen and/or Cre expression on cardiac function. We have previously shown that tamoxifen injection at this dose does not affect basal cardiac function in mice [22].

### 4.2 Embryonic heart isolation

Timed matings were performed, and embryonic age was determined based on the presence of a vaginal plug (E0.5) and confirmed by morphological staging and somite count. Embryonic hearts were microdissected at E14.5 and E17.5 as described previously [12].

### 4.3 Histological analysis

Embryonic and adult hearts were fixed in 4% paraformaldehyde (PFA), dehydrated, embedded in paraffin, and sectioned at 8 µm. Sections were stained with hematoxylin and eosin (H&E) for morphological analysis or Picrosirius Red for collagen deposition according to standard protocols. Left ventricular wall thickness, right ventricular wall thickness, and interventricular septal thickness were quantified at E14.5 and E17.5. Fibrosis area in adult hearts was quantified as the percentage of Picrosirius Red-positive area relative to total myocardial area. All measurements were performed using ImageJ software (NIH) by an investigator blinded to genotype.

### 4.4 Echocardiographic analysis

Transthoracic echocardiography was performed as described previously using a Vevo 2100 ultrasound system equipped with a 32-55 MHz transducer (VisualSonics) [13; 19]. Mice were anesthetized with 1-2% isoflurane and maintained at a heart rate of approximately 500-550 bpm during imaging. For constitutive cKO mice, the same cohort of male mice was evaluated longitudinally at 4, 8, and 12 months of age; female mice were evaluated at 12 months of age. For inducible icKO mice, the same cohort of male mice was evaluated at baseline (pre-injection) and at 2 and 4 months after the final tamoxifen injection. Left ventricular dimensions and function, including ejection fraction (EF), fractional shortening (FS), anterior and posterior wall thickness (LVAWd, LVAWs, LVPWd, LVPWs), and internal dimensions (LVIDd, LVIDs), were measured from M-mode images of the parasternal short-axis view. Diastolic function was assessed in 12-month-old male cKO mice by pulse wave and tissue Doppler echocardiography, and E/E′ and E/A ratios were calculated [20]. All analyses were performed by an investigator blinded to genotype.

### 4.5 Mitochondrial respiration in permeabilized heart tissue

Mitochondrial oxygen consumption was measured in permeabilized left ventricular tissue from 6-month-old male mice using the Oroboros Oxygraph-2k (O2k) system as described previously [13]. We selected this time point to coincide with the exercise capacity assessment. Briefly, oxygen consumption rate (OCR) was measured sequentially under the following conditions: no exogenous substrate (baseline), glutamate (10 mM)/malate (2 mM) to assess complex I-supported leak respiration, ADP (5 mM) to stimulate complex I-driven oxidative phosphorylation, and succinate (10 mM) to engage combined complex I+II-supported respiration. Respiration rates were normalized to tissue wet weight (pmol O /min/mg tissue).

### 4.6 Transmission electron microscopy

Left ventricular tissue from 6-month-old male control and Perm1cKO mice was fixed in 2.5% glutaraldehyde in 0.1 M cacodylate buffer, post-fixed in 1% osmium tetroxide, dehydrated through a graded ethanol series, and embedded in epoxy resin. Ultrathin sections (60-80 nm) were stained with uranyl acetate and lead citrate and imaged using a transmission electron microscope [20]. For each animal, non-overlapping fields, which contain the sarcolemma, were acquired from longitudinally oriented ventricular tissue. Mitochondria were classified by ultrastructural location [39; 47]: subsarcolemmal mitochondria (SSM) as those located between the sarcolemma and the outermost myofibril, and interfibrillar mitochondria (IFM) as those located between adjacent myofibrils (**Fig. 5F**). Mitochondrial profiles were manually traced in ImageJ [48], and the areas occupied by SSM, IFM, and total mitochondria were each expressed relative to the total tissue area of the field [49]. All quantification was performed by an investigator blinded to genotype.

### 4.7 Protein lysate preparation and Western blotting

Heart and skeletal muscle tissues were homogenized as described previously [9]. Protein concentration was determined using the DC Protein Assay (Bio-Rad). Equal amounts of protein (10 µg per lane) were resolved by SDS-PAGE and transferred to PVDF membranes. Membranes were incubated overnight at 4°C with primary antibodies, followed by HRP-conjugated secondary antibodies. GAPDH served as a loading control unless otherwise stated. Bands were visualized using enhanced chemiluminescence and captured with a ChemiDoc Imaging System (Bio-Rad). Densitometric quantification was performed using ImageJ (NIH) or Image Lab software (Bio-Rad). Antibodies used in this study are listed in Supplemental Table 2.

### 4.8 Quantitative real-time PCR (qRT-PCR)

Total RNA was isolated from heart tissues using TRIzol reagent (Invitrogen) according to the manufacturer’s protocol. cDNA was synthesized from 500 ng of total RNA using a High-Capacity cDNA Reverse Transcription Kit (Thermo Fisher Scientific). qRT-PCR was performed using Power SYBR Green PCR Master Mix (Thermo Fisher Scientific) on an iCycler real-time PCR detection system (Bio-Rad). Gene expression was calculated using the ΔCT method and normalized to the reference gene 36B4 [9]. All primers are listed in Supplemental Table 1.

### 4.9 Mitochondrial DNA copy number

Genomic DNA was isolated from left ventricular tissue [9]. Mitochondrial DNA (mtDNA) copy number was determined by qPCR using primers targeting a mitochondrial gene (CoxII) and a single-copy nuclear gene (Dio3). The ratio of mitochondrial to nuclear DNA was calculated and expressed relative to the control.

### 4.10 Citrate synthase activity and tissue ATP content

Citrate synthase activity was measured as described previously and normalized to total tissue protein [10]. Tissue ATP content was quantified using an ATP Assay Kit (Abcam) according to the manufacturer’s instructions and normalized to total tissue protein. For post-exercise measurements, tissues were collected after a fixed 12-min running protocol (see below) and snap-frozen in liquid nitrogen.

### 4.11 Treadmill exercise testing

Exercise tolerance was assessed using a motorized treadmill (Exer 3/6M, Columbus Instruments) as described previously [50]. Mice were habituated to the treadmill for 3 consecutive days before testing and were given at least 1 day of rest between habituation and the exercise test. Habituation and exercise testing were performed during the daytime between 10:00 AM and 6:00 PM.

For the running-to-exhaustion test, mice ran on a 5% incline starting at 5.0 m/min. For 6-month-old mice, speed was increased in a stepwise manner (5, 10, 15, and 20 m/min) until exhaustion. For 12-month-old mice, speed was increased to 10 m/min and maintained until exhaustion to accommodate age-related differences in baseline running capacity. Mice were motivated to run using bristled brushes at the back of the treadmill and were considered exhausted when they could no longer run in response to this stimulus [51]. Running time and total distance were recorded.

For post-exercise metabolic analysis, a separate cohort of 6-month-old mice underwent a fixed 12-min running protocol under the exhaustion test conditions described above (5 m/min 5 min → 10 m/min 5 min → 15 m/min 2 min). This time point was selected to precede exhaustion in cKO mice, enabling comparison of cardiac energetics at a matched submaximal workload. Immediately after the 12-min run, we collected plasma and harvested hearts and gastrocnemius muscles for lactate and ATP measurements. Plasma lactate levels were assessed as described previously [50]. Sedentary controls from the same cohort were handled identically but did not run on a treadmill.

### 4.12 Isolation and culture of neonatal mouse cardiomyocytes and adenovirus infection

Primary mouse neonatal ventricular cardiomyocytes (NCM) were isolated from postnatal day 0 to 2 (P0 to P2) C57BL/6 mice [12; 52]. Briefly, newborn pups were decapitated, and ventricles were excised, cut into eight pieces, and washed in ice-cold HBSS without Ca^2+^ and Mg^2+^. After washing, hearts were placed in 20 ml HBSS containing trypsin and incubated for 4 hr at 4°C. Hearts were digested with HBSS containing collagenase at 37°C. Isolated cells were plated in non-coated plastic dishes with NCM culture media [30% DMEM, 30% M199, 25% FBS, and 15% horse serum] and cultured for 2 hrs at 37°C (termed pre-plating) to remove fibroblasts and endothelial cells. After pre-plating, non-adherent cardiomyocytes were seeded at 5x10^5 cells/well in 12-well plates. After 24 hr incubation, cells were washed with PBS to remove dead cells and debris. For the experiments described, NCM were infected with adenoviruses expressing shGFP (Control) or shPerm1 at a multiplicity of infection (M.O.I.) of 50 [9]. After 48 h of infection, cells were subjected to further assays as discussed.

### 4.13 β-Adrenergic stimulation and cellular ATP measurement

Prior to stimulation, cardiomyocytes were serum-starved for 1 h in DMEM. Isoproterenol hydrochloride (Sigma-Aldrich) was dissolved in PBS immediately before use and added to cells at a final concentration of 1 µM for 30 min at 37°C; vehicle-treated cells received an equal volume of PBS.

After treatment, cells were lysed, and total ATP content was quantified using an ATP Assay Kit (Abcam) according to the manufacturer’s instructions. Luminescence values were confirmed to fall within the linear range of a freshly prepared ATP standard curve. ATP content was normalized to total cellular protein determined by DC Protein Assay (Bio-Rad) from parallel wells, and expressed relative to vehicle-treated shGFP cells. All four experimental groups were processed on the same plate on the same day. Data are from four independent biological replicates.

### 4.14 Statistical analysis

Data are presented as mean ± SEM. Comparisons between two groups were performed using unpaired two-tailed Student’s *t-tests*. For echocardiographic data, two-way ANOVA was used with genotype and time as factors, followed by Bonferroni’s multiple comparisons test. For experiments with two independent variables and independent groups, including genotype × exercise condition for tissue ATP and plasma lactate measurements and genotype × isoproterenol treatment for cellular ATP measurements, we used two-way ANOVA followed by Bonferroni’s multiple comparisons test. Normality of residuals and homogeneity of variance were confirmed by Shapiro-Wilk and Levene’s tests, respectively. A value of *P* < 0.05 was considered statistically significant. All statistical analyses were performed in GraphPad Prism.

## CRediT

Y.C. designed experiments, performed experiments, analyzed data, and wrote the manuscript. R.S.R. was involved in experimental planning, and data analysis on some of the initial studies and reviewed and edited the manuscript. S.T., T.L., and W.F. performed experiments and analyzed data. S.T., S.S., J.C., R.S.R., S.L., and Y.C. reviewed and edited the manuscript.

## Acknowledgements

We thank all lab members for their technical assistance. We thank Dr. Anastasia Kralli for her advice on experimental planning and data discussion.

## Data Availability

Data are available upon reasonable request.

---

**Supplemental Figure S1. Cardiomyocyte-specific Perm1 deletion preserves basal cardiac structure and function in female mice at 12 months of age.** Cardiac phenotype was assessed by transthoracic echocardiography in 12-month-old female control (CTL) and Perm1cKO mice. (**A**) Systolic function, including heart rate (HR), ejection fraction (EF), and fractional shortening (FS). (**B-D**) Echocardiographic structural parameters, including left ventricular anterior and posterior wall thickness in diastole and systole (LVAWd, LVAWs, LVPWd, LVPWs) and left ventricular internal dimensions in diastole and systole (LVIDd, LVIDs). Data are mean ± SEM; n = 6 per genotype. Statistical analysis was performed using an unpaired two-tailed Student’s t-test. N.S., not significant.

**Supplemental Figure S2. Diastolic function is preserved in Perm1cKO hearts at 12 months of age.** Diastolic function was assessed by pulse wave and tissue Doppler echocardiography in 12-month-old male control (CTL) and Perm1cKO mice. (**A**) Ratio of early to late transmitral filling velocity (E/A). (**B**) Ratio of early transmitral filling velocity to early mitral annular velocity (E/E′). Data are mean ± SEM; n = 5 per genotype. Statistical analysis was performed using an unpaired two-tailed Student’s t-test. N.S., not significant.

**Supplemental Figure S3. Validation of Perm1 knockdown in neonatal mouse cardiomyocytes.** Neonatal mouse cardiomyocytes were transduced with adenovirus expressing shRNA targeting Perm1 (shPerm1) or GFP (shGFP; control) at an MOI of 50 and harvested 48 h after infection. Perm1 mRNA was quantified by RT-qPCR, normalized to 36B4, and expressed relative to shGFP. Perm1 expression was reduced by approximately 80% in shPerm1-transduced cells. Data are mean ± SEM; n = 4 per group. **, P < 0.01. Statistical analysis was performed using an unpaired two-tailed Student’s t-test.

