## Supplementary figures and images for "Cardiomyocyte Perm1 sustains cardiac bioenergetic reserve and exercise capacity"

### Supplemental Figs

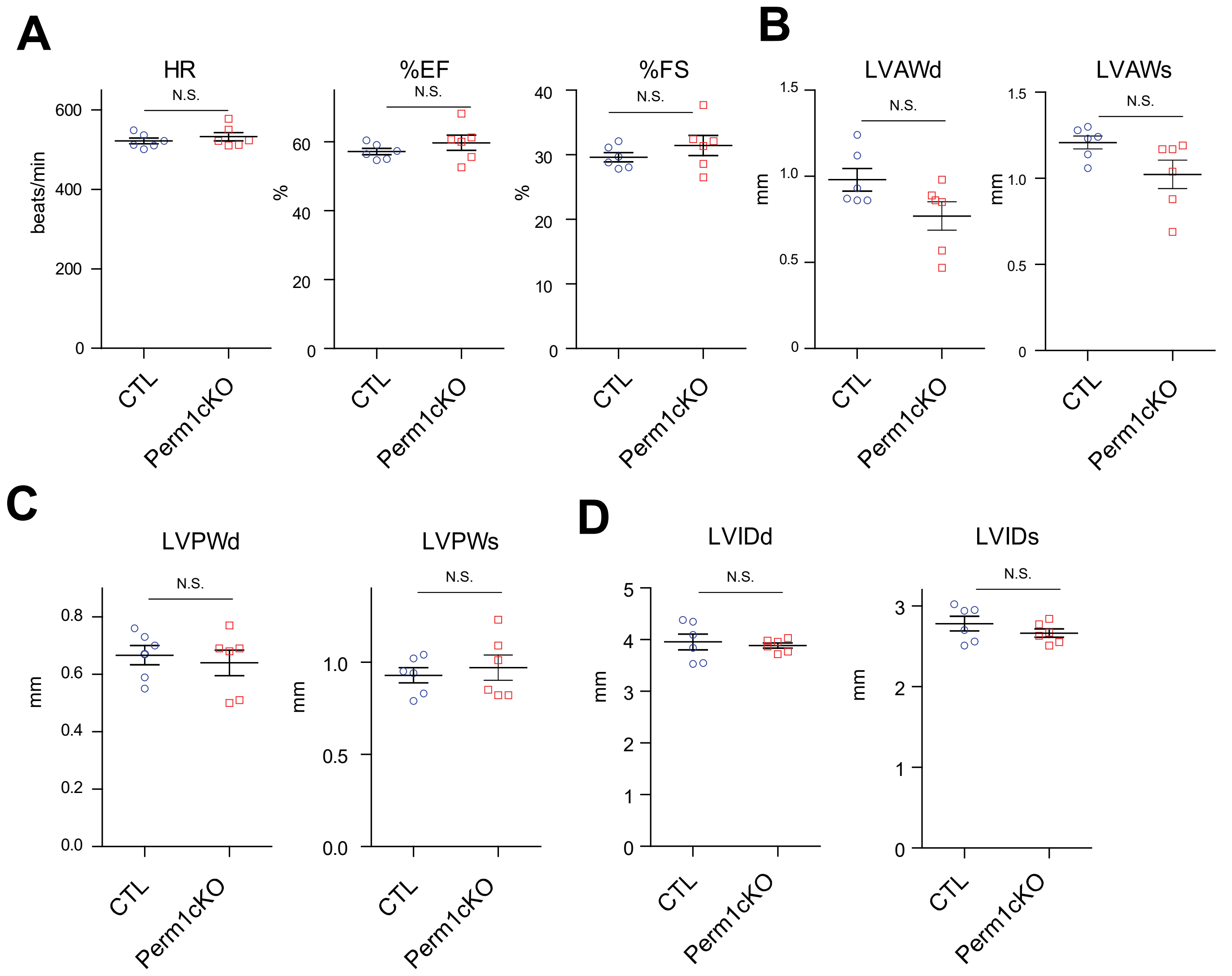


#### Supplemental Fig. S1

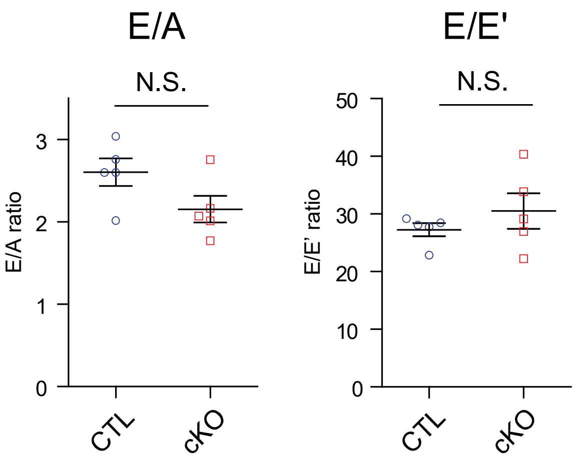


#### Supplemental Fig. S2

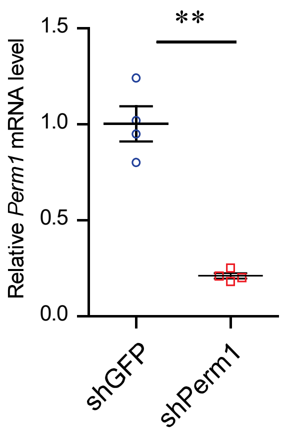


#### Supplemental Fig. S3
