## Supplemental table 1 for "Cardiomyocyte Perm1 sustains cardiac bioenergetic reserve and exercise capacity"

| Supplemental Table 1. Oligonucleotide sequences used in this study. | | |
|---|---|---|
| **Name** | | |
| **CRISPR/Cas9 reagents for generating the Perm1 floxed allele** | | |
| Perm1-L crRNA | AGAUAGGAGGACAGACCCUGGUUUUAG AGCUAUGCUGUUUUG | |
| Perm1-R crRNA | CACUGUCCUAGAUGCCAUCAGUUUUAG AGCUAUGCUGUUUUG | |
| Perm1-L donor ssDNA | ATTGGGCAAAATGGGGCCTTGGTTAGC AGAGCAGGAATAGATAGGAGGACAGAC CATAACTTCGTATAGCATACATTATAC GAAGTTATCTGAGGACAGCCAGAATGG ACTGCTGGAGTTCAGGGAGTTCTTATT TAAGTTCTA | |
| Perm1-R donor ssDNA | AAGCCAGACACTTCAGGGGTTTGGGCC TTGTCACTGACCACTGTCCTAGATGCC AATAACTTCGTATAGCATACATTATAC GAAGTTATTCAGGGACTTTCCTCTAGC TTTTAAGGGCCTGGCACTAGAGGCCCA CTCACACTC | |
| **Name** | **Forward / Sense (5'→3')** | **Reverse / Antisense (5'→3')** |
| **Genotyping PCR primers** | | |
| Perm1 floxed allele (5' loxP) | GAGAGGAGAGAAAGGTTGATAGG | GAATCTATAGTGTCCAGCAGGTGA |
| Perm1 floxed allele (3' loxP) | AAGAATAATCTCCCACGCTATC | AGAGATCTTTTTCTCACACTG |
| Cre transgene | TGCCTGCATTACCGGTCGATGC | CCATGAGTGAACGAACCTGGTCG |
| **Quantitative RT-PCR primers** | | |
| Perm1 | TGCCTATTTCCATCCCTGAG | GGCACATGTCGTTTTGACTG |
| Pgc-1α (Ppargc1a) | CCTCATAAAGCCAACCAAGATAA | GTTTAGTCTTCCTTTCCTCGTGTC |
| Pgc-1β (Ppargc1b) | CTCCTCCTGGCCACATCCT | ACACAAGGCCGCTGACTTC |
| Errα (Esrra) | GGAGGACGGCAGAAGTACAAA | GCGACACCAGAGCGTTCAC |
| Errβ (Esrrb) | GGAATGCTAAAACGGGACTGA | TGATGAAGGAGCCGCAACTA |
| Errγ (Esrrg) | TCCCCGACAGTGACATCAAA | GTGTGGAGAAGCCTGGAATA |
| Pparα (Ppara) | AAGGCTATCCCAGGCTTTGC | TTTAGAAGGCCAGGCCGATCTC |
| Pparδ (Ppard) | GTATGCGCATGGGACTCAC | GTCTGAGCGCAGATGGACT |
| Tfam | caaaggatgattcggctcag | aagctgaatatatgcctgcttttc |
| Nrf-1 | CCACGTTGGATGAGTACACG | CTGAGCCTGGGTCATTTTGT |
| Nrf-2 (Gabpa) | CCGCTACACCGACTACGATT | ACCTTCATCACCAACCCAAG |
| Mfn2 | GAAAGTCGAGGCTCTGGATTCA | GACTGGGTGCTTTCCTTCTCTCT |
| Glut4 (Slc2a4) | GATTCTGCTGCCCTTCTGTCC | CGCTCTCTCTCCAACTTCCG |
| Ldhb | CCAGCAGGAGGGGGAGAGT | TGCTTAGGTAGCCCGCTCAG |
| Pdk4 | GTTCCTTCACACCTTCACCAC | CCTCCTCGGTCAGAAATCTTG |
| Lpl | CCTGATGACGCTGATTTTGTAGA | CAGCAGGGAGTCAATGAAGAGA |
| Fabp3 | AAGTGGAACGGGCAGGAGA | GAGGAGCGGGCGGTCAG |
| Cpt1b | TCTAGGCAATGCCGTTCAC | GAGCACATGGGCACCATAC |
| Lcad (Acadl) | TCTTTTCCTCGGAGCATGACA | GACCTCTCTACTCACTTCTCCAG |
| Atgl | CCTGATGACCACCCTTTCCAAC | ACCCGTCTGCTCTTTCATCCAC |

| | | |
|---|---|---|
| Ckmt2 | TGTCTGAGATGACGGAGCAG | CCTGGTGTGGTCTTCCTCAT |
| Idh3a | AGGACTGATTGGAGGTCTTGG | ATCACAGCACTAAGCAGGAGG |
| Cox7a1 | GACAATGACCTCCCAGTACAC | GCCCAGCCCAAGCAGTATAAG |
| Anf (Nppa) | TACAGTGCGGTGTCCAACACAG | TGCTTCCTCAGTCTGCTCACTC |
| Bnp (Nppb) | CTGAAGGTGCTGTCCCAGAT | CCTTGGTCCTTCAAGAGCTG |
| 36B4 (Rplp0) | CTGTGCCAGCTCAGAACACTG | TGATCAGCCCGAAGGAGAAG |
| **Primers for mitochondrial DNA** | | |
| mt-CoxII (mt-Co2) | TCTCCCCTCTCTACGCATTCTA | ACGGATTGGAAGTTCTATTGGC |
| Dio3 | GGAGGTTGTCCGACCTGAT | GTCCCTTGTGCGTAGTCGAG |
