## Supplemental table 2 for "Cardiomyocyte Perm1 sustains cardiac bioenergetic reserve and exercise capacity"

**Supplemental Table 2. Antibodies used in this study.**

| Target | Source | Catalog No. |
|---|---|---|
| **Target protein** | | |
| PERM1 | Sigma-Aldrich | HPA031711 |
| **MICOS complex and mitochondrial dynamics** | | |
| Mitofilin / IMMT (MIC60) | Cell Signaling Technology | 30671 |
| CHCHD3 (MIC19) | Novus Biologicals | NBP1-83656 |
| CHCHD6 (MIC25) | Novus Biologicals | NBP1-91779 |
| DRP1 | Cell Signaling Technology | 8570 |
| OPA1 | Cell Signaling Technology | 80471 |
| **Oxidative phosphorylation** | | |
| Total OXPHOS rodent cocktail (CI–CV) | Abcam | ab110413 |
| SDHA (Complex II) | Cell Signaling Technology | 11998 |
| **Metabolic enzymes** | | |
| PDH (pyruvate dehydrogenase) | Cell Signaling Technology | 2784 |
| PDK4 | Nuvos Biologicals | NBP1-07047 |
| CKMT2 | Proteintech | 13207-1-AP |
| Hexokinase I (HKI) | Cell Signaling Technology | 2024 |
| Hexokinase II (HKII) | Cell Signaling Technology | 2867 |
| CPT1b | Cell Signaling Technology | 41803 |
| LCAD (ACADL) | Proteintech | 17442-1-AP |
| ATGL (PNPLA2) | Cell Signaling Technology | 2138 |
| **Loading control** | | |
| GAPDH | Cell Signaling Technology | 2118 |
| **Secondary antibodies** | | |
| Anti-rabbit IgG, HRP-linked | Cell Signaling Technology | 7074 |
| Anti-mouse IgG, HRP-linked | Cell Signaling Technology | 7076 |
